# The role of clearance in neurodegenerative diseases

**DOI:** 10.1101/2022.03.31.486533

**Authors:** Georgia S. Brennan, Travis B. Thompson, Hadrien Oliveri, Marie E. Rognes, Alain Goriely

## Abstract

Alzheimer’s disease, the most common form of dementia, is a systemic neurological disorder associated with the formation of toxic, pathological aggregates of proteins within the brain that lead to severe cognitive decline, and eventually, death. In normal physiological conditions, the brain rids itself of toxic proteins using various clearance mechanisms. The efficacy of brain clearance can be adversely affected by the presence of toxic proteins and is also known to decline with age. Motivated by recent findings, such as the connection between brain cerebrospinal fluid clearance and sleep, we propose a mathematical model coupling the progression of toxic proteins over the brain’s structural network and protein clearance. The model is used to study the interplay between clearance in the brain, toxic seeding, brain network connectivity, aging, and progression in neurodegenerative diseases such as Alzheimer’s disease. Our findings provide a theoretical framework for the growing body of medical research showing that clearance plays an important role in the etiology, progression and treatment of Alzheimer’s disease.

## 1 Introduction

Neurodegenerative diseases such as Alzheimer’s disease (AD) are progressive disorders associated with a gradual loss of cognitive faculties and, ultimately, a state of dementia. These diseases are characterized by the presence of proteins, present under healthy conditions, that misfold, propagate and replicate in the pathological state. The study of neurodegenerative diseases in humans is a formidable challenge; investigating fundamental neurological questions often requires invasive experimental techniques. Advances in imaging technology, and mechanisms discovered using animal models, now allow for the construction of brain graphs, from patient data, and dynamical systems modeling neurodegenerative disease progression in humans. In this manuscript, we propose a novel network neurodegeneration model that couples together the spreading, replication and dynamic removal of misfolded proteins within the brain.

### 1.1 Prions and the prion-like hypothesis of neurodegeneration

The term *prion* refers to a type of protein that can trigger normal proteins to fold abnormally. The use of the word ‘prion’ originates [89] in the seminal work of Stanley Prusiner, for which he received a Nobel prize, on transmissible spongiform encephalopathies (TSE). The British scientist J.S. Griffith had previously speculated that the causative agent of scrapie, a TSE in sheep, was proteinaceous in nature. Prusiner isolated the infectious proteinaceous agent and showed that it could only be inactivated using methods for destroying proteins [89], thus settling a prominent debate at the time. It was shown that scrapie resulted from the naturally occurring cellular prion protein (PrP^C^) being transformed to a toxic, misfolded confirmation (PrP^SC^) and that otherwise healthy PrP^C^ protein would misfold into PrP^SC^ in the presence of existing PrP^SC^.

The autocatalytic prion replication mechanism is illustrated in Figure 1. A healthy (blue) and misfolded (red) version of a protein become proximal (blue/red joined), misfolding is transmitted (red/red joined) and the result is two copies of the misfolded protein. The *prion-like hypothesis* of neurodegenerative diseases posits that the misfolded proteins that characterize many human neurodegenerative pathologies (Section 1.2) spread their misfolded state as prions do. One could argue that the prion-like hypothesis also originated with Prusiner [59] but, regardless of its origin, recent evidence for this hypothesis has been mounting steadily [13, 26, 38, 39, 45, 52] across neurodegenerative pathologies and the basic mechanism of prion-like self replication in neurodegenerative diseases is not controversial.

**Figure 1:**
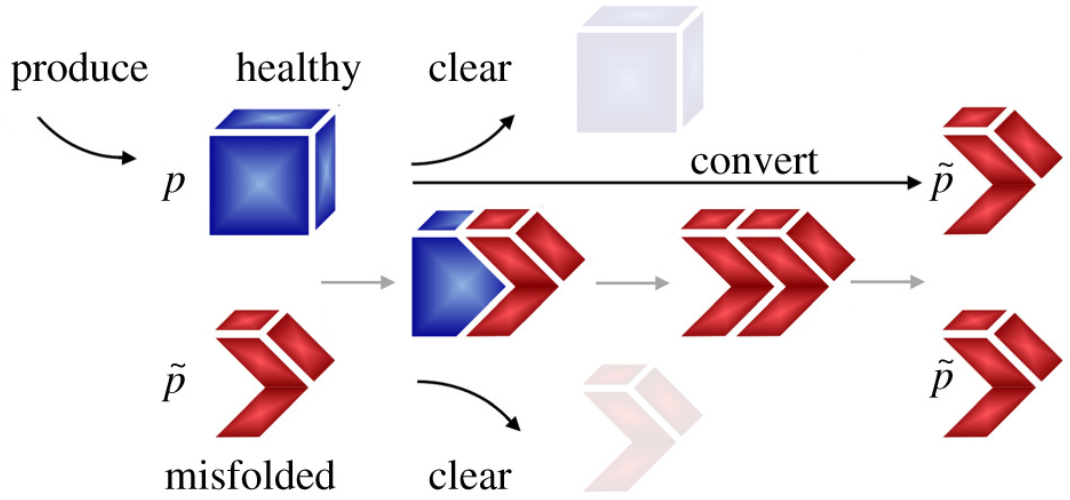
A healthy protein (blue) is converted to a misfolded protein (red) by the autocatalytic prion, and prion-like, process. Healthy and misfolded toxic proteins can also be removed by brain clearance mechanisms (transparent blue, red).

### 1.2 Typical proteins involved in neurodegenerative diseases

The most common neurodegenerative disease is AD, involving two main proteins: amyloid-*β* (A*β*) and tau protein (τP). A*β* results from the cleavage of amyloid precursor protein (APP), a membrane protein highly expressed in neurons and involved in many beneficial brain functions [28]. If an APP is first processed by an enzyme called *α*-secretase, the end result is not harmful. An APP processed instead by an enzyme called *β*-secretase results in toxic, misfolded A*β*. Tau is a protein mostly found in neurons and acts to stabilize microtubules, which provide the physical infrastructure for the delivery of resources from the cell body down the axons. The prevailing hypothesis is that normal tau protein that becomes hyperphosphorylated loses its physiological function and forms toxic, misfolded τP [51]. The next most common neurodegenerative disease is Lewy body dementia and the associated protein is *α*-synuclein which, in its healthy form, helps to regulate the mitochondria of neurons. A likely cause for the transition from normal to misfolded *α*-synuclein is not currently known, but it has been shown that increased temperature and decreased pH favor the appearance of toxic, misfolded *α*-synuclein [5]. Misfolded A*β*, τP and *α*-synuclein all propagate their toxic misfolded conformation in a prion-like fashion [26].

Other, less common, neurodegenerative diseases involve different proteins but the general idea remains the same: a particular brain protein serves a physiological role but, under some pathological conditions, becomes misfolded, is toxic to the brain and propagates the misfolding in a prion-like manner (Figure 1). Moreover, these misfolded proteins, if not removed by the brain (Section 1.4), aggregate into various structures. Single misfolded proteins, or small oligomeric structures, that are prone to causing prion-like templated misfolding (Section 1.1) and further aggregation are called (toxic or misfolded) *seeds*. Toxic seeds are motile and can propagate from one brain region to another (Section 1.3); large, aggregated protein structures form immobile aggregated deposits. Aggregated deposits of misfolded protein fragments that build up in the cellular space outside of neurons are referred to as *plaques* while aggregated twisted-strand strand structures that form inside of neurons are called *neurofibrillary tangles* (NFT).

### 1.3 Regional transmission of misfolded proteins in the brain

Toxic, misfolded seed proteins are generally motile and can propagate between axonally-connected brain regions [17, 83]. Regional seed propagation has been noted in humans [6, 38] and animal models [12]. In vitro evidence demonstrated the direct neuronal exchange of misfolded seed proteins [49] using axonally-related pathways; further studies [87] show that seeds can move bidirectionally, along axons, and are trafficked between synaptically connected neighbors. Additional experiments have observed that seeds can also make their way into the extracellular space [46, 56, 58] where they would then be subjected to strongly axonally-anisotropic extracellular fluid diffusion.

### 1.4 Removal systems for brain waste and misfolded proteins

The brain is a dynamic, metabolically-active organ; errors in protein synthesis and folding occur relatively frequently and the brain has several *clearance systems* that remove brain waste products, including misfolded and unnecessary, but otherwise healthy, proteins. Brain waste clearance systems are, broadly speaking, based on the blood circulation, based on cellular recycling or based on the circulation of the water-like cerebrospinal fluid [74, 50]. Molecular chaperones allow some waste material to cross over the blood-brain barrier and be carried out of the brain through the blood circulation. Cellular recycling can happen either extracellularly, such as the degradation of waste proteins by proteases, or intracellularly, via the ubiquitin-proteasome, autophagy-lysosome or endosome-lysosome pathways. The cerebrospinal fluid circulation is now thought to play an important role in brain clearance; circulating cerebrospinal and interstitial fluid can carry waste proteins to nearby lymphatic vessels that reside outside the brain parenchyma [50, 66].

Brain clearance systems have been proposed to play a potential role in the etiology and progression (Section 1.5) of neurodegenerative diseases but the precise mechanisms, and how they are altered in neurodegenerative diseases, are topics of ongoing research [31, 32, 50, 72, 74]. Moreover, the efficacy of the brain’s various clearance systems appear to decline in the presence of toxic, misfolded proteins [4, 9, 10, 31, 50] and this decline may lead to the large-scale proliferation of misfolded brain proteins. The combined action of brain clearance systems is represented as the world ‘clear’ in Figure 1, alongside the removed healthy (transparent blue) and toxic misfolded (transparent red) proteins.

### 1.5 Etiology, progression and subtypes of neurodegenerative diseases

Our current understanding cannot ascribe a singular cause, or distinct set of causes, that must necessarily give rise to neurodegenerative disease. In fact, several hypotheses for the etiology of AD alone have been advanced and include: the A*β* hypothesis; the tau hypothesis; the lymphatic hypothesis; the vascular hypothesis; and the notion of a combination of genetic and lifestyle factors [20, 41], among others. Whether there is a singular set of pathogenic factors or if neurodegeneration emerges from the multifactorial interactions of system failures is currently an open question.

Despite a lack of clear etiological factors for neurodegenerative diseases, many neurodegenerative diseases exhibit *temporal staging patterns* that can act as a measure of disease progression. A temporal staging pattern is defined by a relatively structured ordering in which protein plaque or NFT aggregates (Section 1.2) appear across disease-specific regions throughout the brain [6, 11, 38, 39]. Aggregates are thought to accumulate following the regional appearance of a toxic seed (Section 1.3), either locally or from an axonally connected region [17, 83]. In AD, for instance, τP aggregates are thought to follow a hierarchical staging pattern referred to, generally, as a patient’s Braak stage [6, 11], demarcating their severity of AD progression (Figure 2). Experiments suggest [6, 11] that aggregated τP NFT deposits appear first in the entorhinal cortex (Figure 2 left, purple) of AD patients and subsequently appear in stages II through V/VI as AD progresses while τP NFT continue to accumulate in the regions defining the previous stages; other neurodegenerative diseases show similar, but disease-specific, regional progression patterns [38].

**Figure 2:**
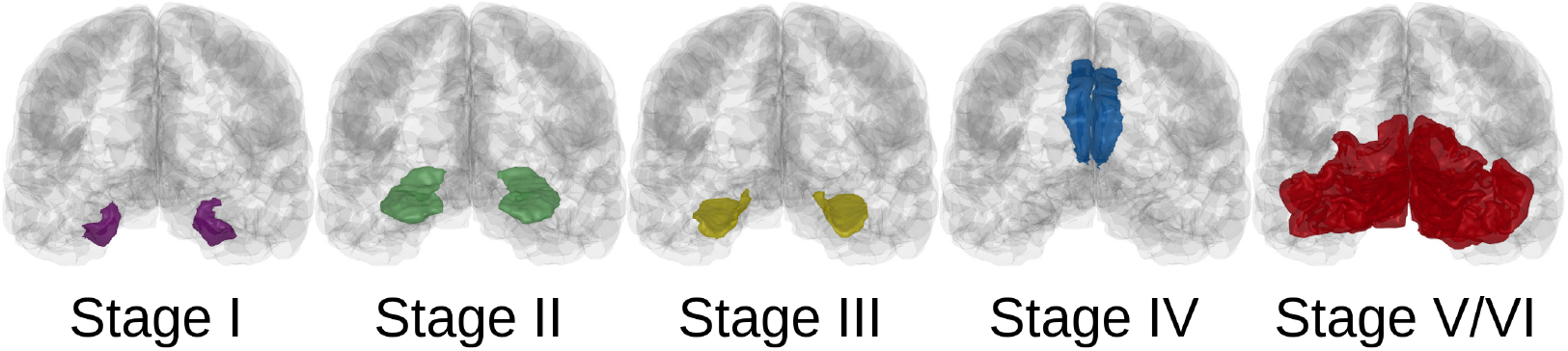
The Braak staging pattern of τP NFT deposition in Alzheimer’s disease (adapted from [17]); posterior coronal view of a glass brain. Stage I: entorhinal cortex (purple), Stage II: hippocampus (green), Stage III: parahippocampal gyrus (yellow), Stage IV: rostral and caudal anterior cingulate (blue), Stage V/VI: cuneus, pericalcarine cortex, lateral occipital cortex, lingual gyrus (red).

The canonical gold standard for confirming a neurodegenerative disease diagnosis, and disease stage, is a postmortem analysis [23] of the regional distribution of aggregated (plaque or NFT) brain proteins. In AD, regional variations in the predominance of τP NFT aggregate deposition, traditionally determined postmortem, have been linked to different *subtypes* [23, 37, 47, 81] that manifest differences in AD clinical presentation including the rate of cognitive decline, disease duration and age of onset [37, 47].

### 1.6 Mathematical models of network neurodegeneration

Due to difficulties in assessing pathology in vivo, mathematical modeling is now being hailed by the medical community as a powerful new research tool to improve the understanding of neurodegenerative diseases [43, 48, 77]. The construction and analysis of the topological properties of brain networks, derived from patient imaging data, dates back to at least the turn of the century [7, 73]. The study of brain network structure has provided invaluable insights into regional and functional connectivity and has elucidated how network properties are altered as a result of neurodegeneration. However, these types of mathematical approaches do not speak to the dynamics of neurodegenerative diseases.

Dynamical models of neurodegenerative diseases should reflect the central mechanisms, suggested by clinical experiment, thought to drive the progression of pathology. The most fundamental mechanisms suggested in the literature are now the prion-like hypothesis (Section 1.1), of autocatalytic misfolded protein replication, and the transmission of misfolded brain proteins (Section 1.3) from one brain region to its axonally-connected neighbors. Some of the first protein pathology models [53, 62, 63] employed a structural brain network to evolve the local misfolded protein concentration using a linear system of differential equations based on the network’s graph Laplacian matrix. This early *network diffusion modeling* had the advantage of an analytic solution and managed to recapture some correlation with neuroimaging data for various dementias.

The conception of the network diffusion modeling predated-dated the widespread resurgence of the prion-like hypothesis in the literature. As a result, the primary drawback of these models was that they did not include autocatalytic replication, nor other contemporary mechanistic hypotheses, and were understandably limited in their ability to explain patient-specific neuroimaging data. The prion-like hypothesis has now been incorporated into more sophisticated *network neurodegeneration models* that include both autocatalytic replication and weighted graph Laplacian axonal propagation [24, 25] in the context of diffusion-reaction systems posed on structural brain network graphs. Early network neurodegeneration models have been used to predict patient-specific disease trajectories [68, 67, 69] and more recent extensions have been used to model the experimentally observed phenomenon of interaction between A*β* and τP proteins in Alzheimer’s disease [75]. However, previous network neurodegeneration models have failed to account for regional heterogeneity in clearance, the adverse effects (Section 1.4) of toxic proteins on brain clearance, and the potential for the dynamic interplay between toxic proteins and brain clearance to influence the etiology and progression (Section 1.5) of neurodegenerative diseases.

In this manuscript, we advance a novel network neurodegeneration model to study the role of a dynamically evolving clearance (Section 1.4) in neurodegenerative diseases. The model couples a reduced-order toxic propagation equation, derived from a Smoluchowski system of oligomer aggregation [76], with a first-order evolution equation accounting for the adverse effects of toxic proteins on brain clearance systems. We will see that clearance confers a topologically distributed resilience against neurodegeneration, that clearance efficacy delays disease onset and that regional heterogeneity in clearance can potentially explain the manifestation of primary Alzheimer’s disease subtypes. Overall, our work suggests that variations in clearance may play a key role in the formation and propagation of neurodegenerative pathology.

## 2 Modeling toxic protein and clearance interplay

Network neurodegeneration models are motivated by two primary observations. First, the prionlike hypothesis of neurodegeneration postulates a local, autocatalytic reproduction mechanism for misfolded proteins (Section 1.1). Second, the region-to-region propagation of toxic proteins takes place via the axonal bundles connecting brain regions (Section 1.3). In this section, we describe a diffusion-reaction model posed a network graph. The model captures the regional transmission of misfolded proteins via a weighted graph Laplacian diffusion while the reaction term expresses both local autocatalytic misfolded protein replication in addition to a novel accounting for the effects of toxic proteins on the efficacy of the brain’s clearance systems (Section 1.4).

### 2.1 Network models of toxic protein propagation

To model experimentally observed dynamics of misfolded neurodegenerative protein, we consider a concentration *p* of this toxic protein that diffuses between brain regions and whose local concentration is governed by clearance and an autocatalytic replication that further mediates clearance efficacy. We assume that the protein distribution evolves on the connectome. A structural connectome is a network *G* = (*V, E*), whose node set *V* contains *N* nodes corresponding to brain regions of interest (ROIs) and whose edge set *E* represents axonal fibers between these regions. Given a structural connectome *G*, *p_i_* = *p_i_*(*t*) denotes the toxic protein concentration associated with the node *i* at time *t* and represents the average concentration in the corresponding ROI. A general network neurodegeneration model including toxic protein transport and local clearance may then take the form [24]: find the concentration *p_i_* for *i* ∈ *V* such that

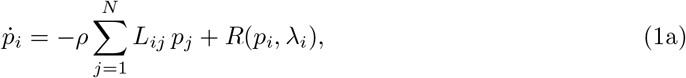

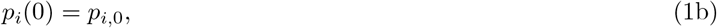

where the superimposed dot denotes the time-derivative, *λ_i_* is the clearance level at node *i*, *L_ij_* are the entries of the graph Laplacian **L** associated with *G*, *ρ* is a transport coefficient, *R* is a reaction relation governing the local dynamics within the ROIs, and *p*_*i*,0_ is the initial value of the concentration on *i*.

The graph Laplacian **L** is defined by **L** = **D** − **W** where **W** is a weighted adjacency matrix of *G* and **D** is the diagonal matrix **D** = diag(*d*_1_,…, *d_N_*), where *d_i_* is the weighted degree associated with node *i* defined by

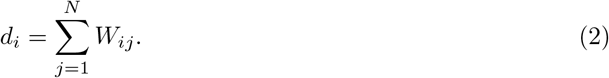

Other normalized forms of the graph Laplacian may and have been used in the literature, but this standard form simultaneously conserves mass and enforces Fick’s constraint, thus guaranteeing that no transport occurs between regions with same concentration [61, Supplementary S1]. Specifically, we select the weighted adjacency matrix with entries 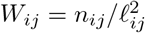, where *n_ij_* is the number of fibers along the axonal tract, connecting node *i* to node *j*, and *ℓ_ij_* is the average fiber length between the same two nodes. The quantities *n_ij_* and *ℓ_ij_* arise from fiber tractography [14] and are provided in the connectome data [40]. This choice of weights is a consistent generalization of conservative finite volume methods applied to diffusion problems [78, 30].

In the remainder of this manuscript, we will use both idealized graphs *G* as well as a human brain connectome with *N* = 1015 nodes (Fig. 3) generated from the data of 426 individual *Human Connectome Project* patients using the Lausanne multi-resolution anatomical parcellation [40, 14] based on the Desikan-Killiany atlas [16].

**Figure 3:**
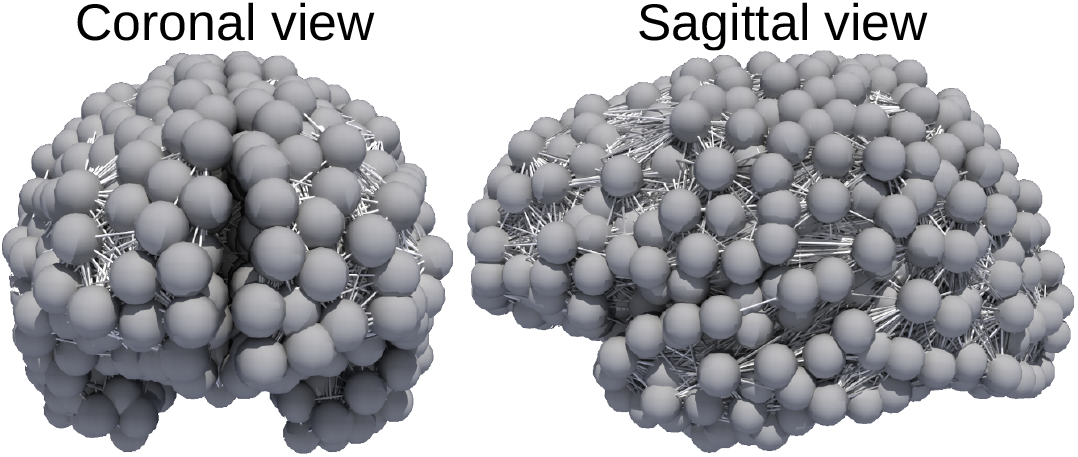
A structural human brain connectome graph. The vertices correspond to a FreeSurfer (Desikan-Killiany) parcellation [15, 16] of the brain’s gray matter. The edges correspond to white matter (axonal) fibers parsed from Human Connectome Project imaging data.

### 2.2 A coupled model of toxic protein transport and brain clearance

Network models can and have been used to investigate various aspects of Alzheimer’s disease, Parkinson’s disease, supranuclear palsy, frontotemporal dementia, and other neurodegenerative diseases. Most early studies consider diffusion w ithout r eaction, i.e. *R* = 0 [62, 1, 63, 54, 84, 55, 53], or use network approaches as a means of interpreting voluminous imaging data sets [34, 82, 81]. Recent work has focused on the autocatalytic nature of protein dynamics leading to a local expansion of the toxic population in agreement with the prion-like hypothesis [25, 75, 61]. In such models, a nonlinear function for *R*(*p_i_*) is chosen to model autocatalytic exponential growth at small concentration and saturation at larger concentration as observed in longitudinal studies [36].

Here, we further generalize network neurodegeneration models to take into account dynamic and heterogeneous brain protein clearance. Brain clearance relies on several different systems (Section 1.4) and may vary in a complex manner from region to region. At the local level, it has recently been shown, using a Smoluchowski model for the dynamics of proteins, that the effect of reducing clearance is to create an instability. Close to this instability, the dynamics takes the universal form of a transcritical bifurcation [76]. Explicitly, continuing to denote by *λ_i_* the level of clearance at node *i*, the normal form of the bifurcation close to *λ_i_* = *λ*_crit_ is

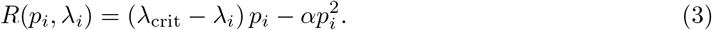

where *α* describes the population expansion and can be obtained from microscopic models [76]. For *λ* > *λ*_crit_ the fixed healthy point *p* = 0 is stable, but it loses stability as *λ* decreases below *λ*_crit_. For *λ* < *λ*_crit_, the fully toxic state instead becomes a stable fixed point. In light of these findings, we define *R* in (1) by (3) and the global evolution of the toxic protein concentration is thus governed by, for *i* = 1,…, *N*:

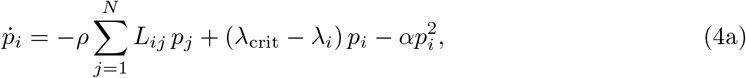

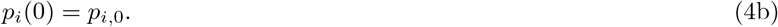

Vice versa, the presence of toxic protein oligomers affects the brain clearance pathways [4, 10, 9, 11, 35, 18, 8, 29]. We model a deteriorating clearance due to the presence of toxic proteins by a first-order rate law:

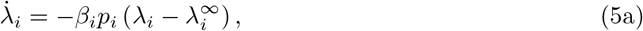

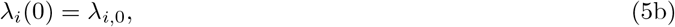

for *i* = 1,…, *N*, where *β_i_* > 0 is a kinetic constant for each *i*, 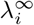 is the minimum regional clearance value. We assume that 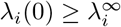. In summary, the full dynamic system is given by: for *i* = 1,…, *N*,

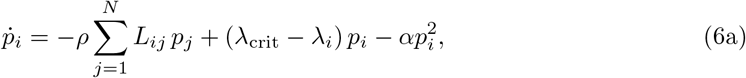

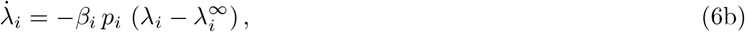

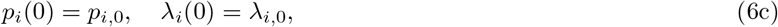

In the case where *λ_i_*(0) = *λ*^∞^ for all *i*, this system reduces to the standard Fisher-Kolmogorov system [24, 85, 61, 68].

## 3 Stability and criticality in the homogeneous coupled model

If the initial conditions and other model parameters are homogeneous i.e. *p*_*i*,0_ = *p*_0_, *λ*_*i*,0_ = *λ*_0_, 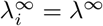, and *β_i_* = *β* for *i* = 1,…, *N*, the system (6) reduces to a homogeneous system equivalent to that of a single node:

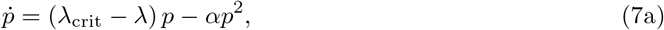

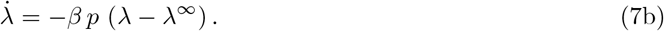

Moreover, in the very early stages of pathology when only a few brain regions have collected any toxic proteins (*p*_*i*,0_ = 0 nearly everywhere and *p*_*j*,0_ ≈ 0 otherwise), the graph Laplacian term in (6a) acts to initiate seeding in nearby neighbors [60] that then evolve according to (7). Thus, advancing a clinical understanding of the stability and fixed points of (7), and their implications, is fundamental to understanding the larger-scale behavior of (6). In Sections 3.1–3.2 we see that (7) admits two fixed points whose clinical significance depends directly on the value of *λ*. In Section 3.3 we will see that (7) admits a notion of criticality that reflects the level of misfolded proteins that the local brain clearance systems can sustain before pathology initiates in the region.

### 3.1 Fixed points and stability

This dynamical system (7) admits a class of fixed points corresponding to the absence of a toxic protein load combined with any clearance *λ*^∞^ ≤ *λ* ≤ *λ*(0):

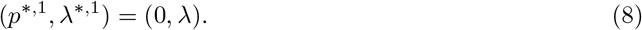

Another fixed point of (7) corresponds to the case in which *p*^*^ > 0 and the clearance is at its minimal value:

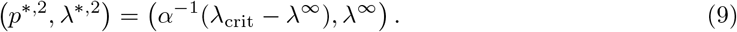

The properties of the Jacobian 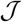 of (7) determine the stability of these fixed points. Specifically, we have that:

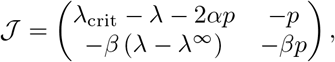

and its eigenvalues are given by

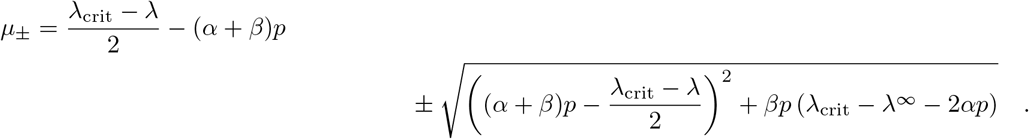

The eigenvalues of 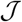 evaluated at (*p*^*,2^, *λ*^*,2^) both have negative real parts; this fixed point is thus unconditionally stable. For the other fixed points, note that the Jacobian’s determinant vanishes at (*p*^*,1^, *λ*^*,1^). However, its local dynamics can be evaluated directly by differentiating (7a) with respect to *p* and evaluating the result at *p*^*,1^ = 0. From these calculations, we note that the fixed point(s) (0, *λ*) are stable if and only if *λ* > *λ*_crit_, and unstable otherwise.

### 3.2 Clinical characterization of fixed points

Combining these fixed points and their stability with their clinical interpretation, we define three regional (nodal) homogeneous disease states: <u>healthy</u>, <u>susceptible</u> and <u>diseased</u> as follows. Each disease state corresponds to fixed points of (7).

I. In the <u>healthy</u> state, (*p, λ*) = (0, *λ*) with *λ* > *λ*_crit_. By (7b), even a small toxic load *p* leads to a reduction in clearance at a rate *β* > 0. However, (7a) dictates that *p* decreases towards to zero as long as *λ* > *λ*_crit_. This state is thus equipped with a degree of resilience to toxic protein seeding and is as such protected from the onset of pathological neurodegeneration.
II. In the <u>susceptible</u> state, (*p, λ*) = (0, *λ*) with *λ* ≤ *λ*_crit_. In this state, clearance is dysfunctional, and a small change in the toxic protein concentration will send the system to the diseased state.
III. In the <u>diseased</u> state: (*p, λ*) = (*α*^−1^(*λ*_crit_ − *λ*^∞^), *λ*^∞^). In this case, the clearance is at its minimal value and the toxic protein concentration is saturated.

The regional characterizations provide a direct perspective on a clinical characterization of the full system (6). First, observe that any region whose state is (9) is, effectively, an incubator of toxic protein seeds. Even if the neighbors of a diseased region are otherwise in the healthy state, (7b) dictates that toxic protein seeds, from the diseased region, will erode the otherwise healthy neighboring clearance values. Toxic seeds will continue to originate in the unstable node, and migrate to adjacent neighbors, until the clearance of the diseased region’s neighbors satisfy *λ* ≤ *λ*_crit_ and, in turn, they transition to diseased regions themselves. Once a node has transitioned to (9), a toxic infection propagates outwards from it and the cascade of deficient clearance, and subsequent invasion, permeates throughout the brain. Thus, we say that the whole brain, as represented by the non-homogeneous system (6), is in the <u>healthy</u> state if all regions satisfy (I), that it is in the <u>susceptible</u> state if at least one region satisfies (II) and that it is in a <u>diseased</u> state if at least one region satisfies (III).

### 3.3 Critical toxic seeding

The term *seeding* refers to the presence of an aggregate prone misfolded protein in a brain region (Section 1.2). If seeding occurs, the seed protein must be cleared by the brain or it will replicate (Section 1.1). Misfolded toxic proteins are thought to be damaging to the brain’s clearance systems (Section 1.4). In fact, the homogeneous system (7) admits a notion of how much toxic protein a region can withstand before the failure of its local clearance capacity. In particular, the healthy and susceptible states differ by whether *λ* > *λ*_crit_ or not. Consider the phase plane in Fig. 4, where orbits for different initial toxic loads *p*_0_ are shown. For *λ*_0_ > *λ*_crit_, each toxic seeding event, yielding *p* > 0, will degrade the clearance capacity until *λ* reaches *λ*_crit_. For sufficiently small seeds, the toxic proteins are cleared and *p* tends to 0. However, for larger seeds, *p* tends to the diseased *p*^*,2^ > 0. Hence, there is a value *p*(0) = *p*_crit_, the *critical toxic seeding*, that sends the system to the susceptible state (0, *λ*_crit_). Conversely, this value is an indicator of the system’s resilience to toxic seeding events.

**Figure 4:**
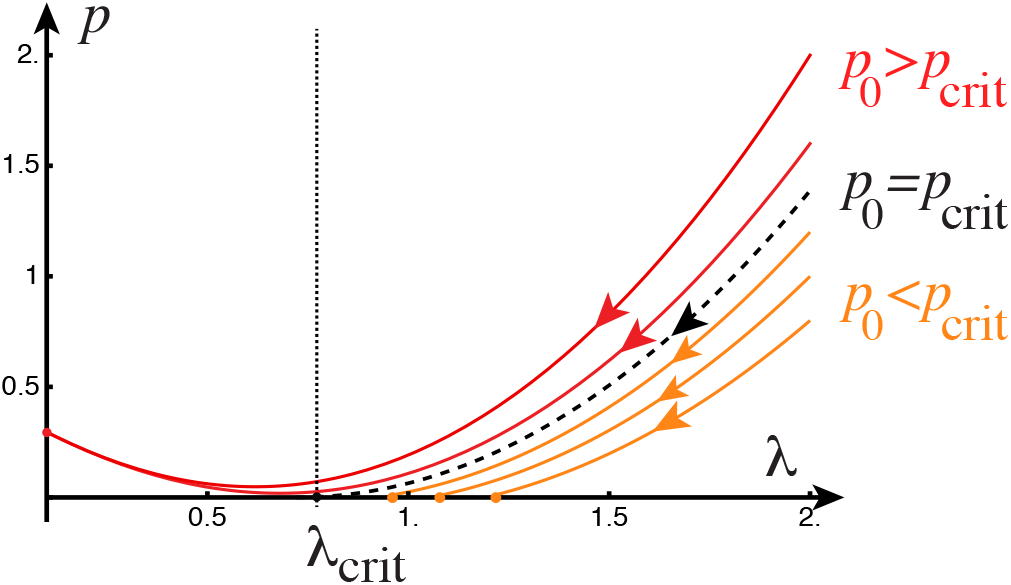
The coupled dynamics of toxic protein evolution versus clearance. An illustrative (*λ, p*) phase plane is shown – with phase paths corresponding to (*λ*_0_, *p*_0_) for *λ*_0_ = 2, and *p*(0) ∈ {0.8, 1.0, 1.2, *p*_crit_, 1.6, 1.8}. The other parameters used are *α* = 2.1, *β* = 1, *λ*_crit_ = 0.72, *λ*^∞^ = 0.1. We find that *p*_crit_ ≈ 1.387. The corresponding diseased state is at *p*^*,2^ ≈ 0.295.

To derive an analytical expression for *p*_crit_ in terms of the model parameters, we examine orbits in the phase plane on the form *p* = *p*(*λ*) and integrate. More precisely, eliminating time derivatives between (7a) and (7b) yield

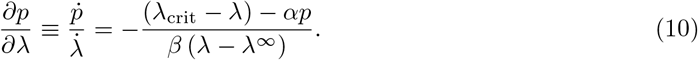

After integrating with respect to *λ* and inserting (*p*(0), *λ*(0)) = (*p*_0_, *λ*_0_), we obtain

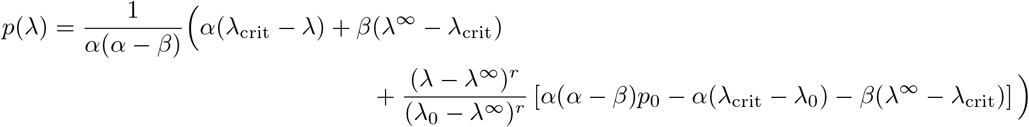

where *r* = *α/β*. The critical toxic seeding is given by the orbit intersecting the *λ*-axis (*p*(*λ*) = 0) at exactly *λ* = *λ*_crit_ with *p*_0_ = *p*_crit_, and is thus given by

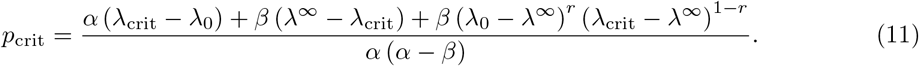

The critical seeding provides important insight into the dynamics of the homogeneous coupled system. Initial seeding values below the critical threshold *p*_0_ < *p*_crit_ will result in a healthy steady state (0, *λ*^*,1^) where *λ*^1,*^ is the largest, strictly positive root of *p*(*λ*) = 0. Conversely, an initial seed of *p*_0_ > *p*_crit_ result in the diseased state.

## 4 Network connectivity increases brain resilience via diffusion and clearance

We now turn from the homogeneous, single-node case to the network case. The analysis of Section 3 demonstrates that clearance effectively contributes to reduce toxic protein load. Next, we will show that a node’s local connectivity can increase its resilience against toxic proteins by relying on the clearance of neighboring regions.

### 4.1 Perturbation analysis of critical network toxic seeding

Consider a node *i* and its neighborhood *G_i_* defined as the set consisting of *i* and the indices of all connected nodes: *j* ∈ *G_i_* iff *L_ij_* ≠ 0. For small *ρ* ≪ 1 (i.e. slow diffusion), the graph Laplacian term in (6) adds a regular perturbation to the homogeneous system (7). Thus, to investigate the effects of connectivity on the critical toxic seeding, we expand the concentration *p_k_* and clearance *λ_k_* for each node *k* in *G_i_* with respect to *ρ*, as

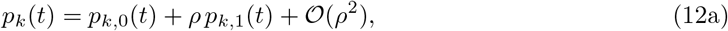

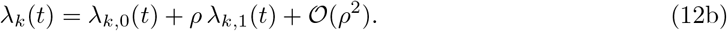

Substituting these into (6), equating powers of *ρ* and dropping the 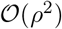 terms yield two sets of equations for each *k* ∈ *G_i_*:

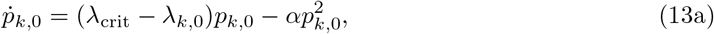

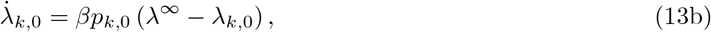

and

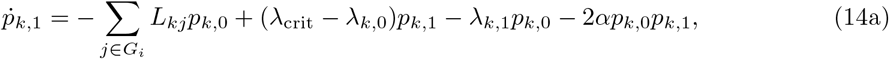

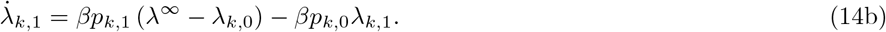

As initial conditions for (13), first consider a toxic seed *p_S_* at node *i* only and a uniform initial clearance *λ*_0_ > *λ*_crit_:

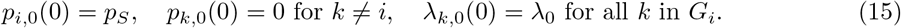

Integrating (13) with (15) allows us to express (14) as a simple inhomogeneous system given by

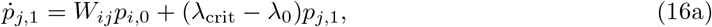

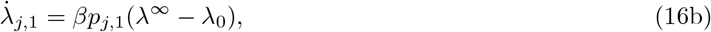

for *j* ≠ *i*. The solution of this equation is, for all nodes *j* in *G_i_* with *i* ≠ *j*,

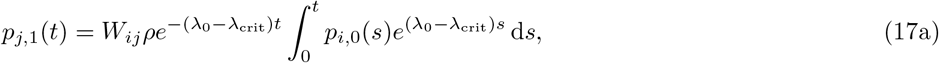

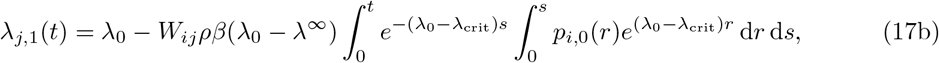

The critical seeding value *p*_crit_, subject to the *ρ* perturbation, is yet to be determined. By definition, if the seeding node is seeded at a level below critical (*p_i_*(0) = *p_S_* < *p*_crit_), then *p_i,_*_0_ decreases, monotonically by (6a), to the asymptotic state 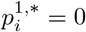. As a result, (17a) implies that

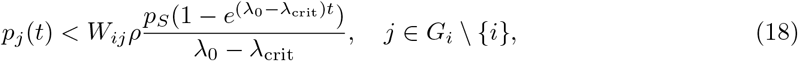

and we conclude that *p_j_*(*t*) < *p_S_* for small perturbations *ρ*. Therefore, when *p_S_* < *p*_crit_, the toxic protein concentration *p_j_* and clearance *λ_j_* in node *j* reach the steady state (0, *λ*^*j*,*^) with *λ*^*j*,*^ > *λ*_crit_.

The task that remains, then, is to ascertain how *p*_crit_ changes as a function of *ρ*. Due to the initial condition (15), the evolution equation (13) gives *p*_*j*,0_(*t*) = 0 for *j* ≠ *i* so that (14a), for *k* = *i*, is given by

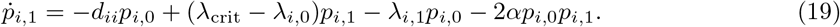

The set of equations (13) and (19) alongside (14b) can now be solved numerically using standard ordinary differential equation solution algorithms, and the perturbed solutions be reconstructed using (12), to quantify the critical network toxic seeding for different connectome configurations.

### 4.2 Critical toxic seeding increases with diffusivity and connectivity

We investigate the impact of alterations in diffusivity and connectivity on the critical network toxic seeding. First, to quantify the effect of diffusion, we estimate the initial condition *p_S_* needed to reach the asymptotic state (0, *λ*_crit_) via the aforementioned numerical procedure, for different *ρ*. For each experiment, we consider a fixed network consisting of one node with five neighbours, let *w_ij_* = 1, *λ*_0_ = 2.0, *α* = 2.1, and *β* = 1, and consider a range of *ρ*. The results demonstrate that the critical network toxic seeding increases with increased diffusivity (Figure 5).

**Figure 5:**
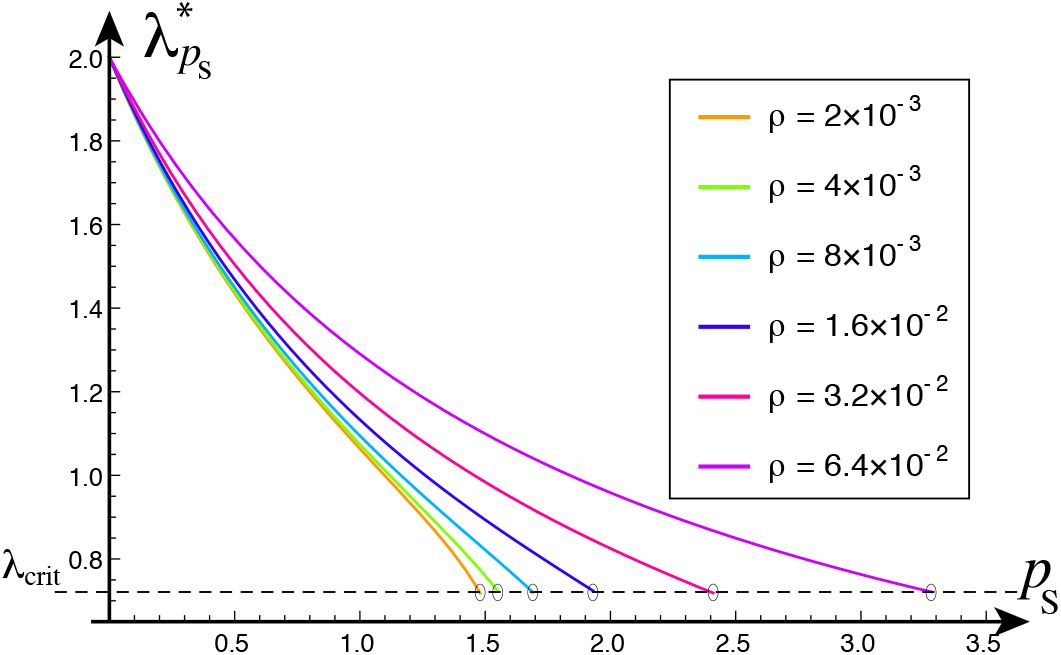
The effect of varying *ρ* on the critical seeding at a node with five neighbors. Plot of clearance *λ* versus toxic seeding *p_S_*. The critical clearance level, *λ*_crit_ = 0.72, is shown as a dashed horizontal line. The final clearance value 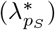 is shown as a function of the seed values (*p_S_*). Critical seed values *p*_crit_ coincide with the intersection of 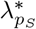 with the dashed line (circles).

Second, we are interested in the effect of brain connectivity on the critical toxic network seeding. Letting the node degree measure regional brain connectivity, we consider a series of numerical experiments with a node *i* and increasing node degree *d_i_* i.e. an increasing number of connected neighbours. Again, we observe that the critical toxic network seeding *p*_crit_ increases with the degree (Figure 6). This analysis suggests that the brain’s connectivity may protect regions by allowing them to share the burden of clearing toxic loads with their neighbors.

**Figure 6:**
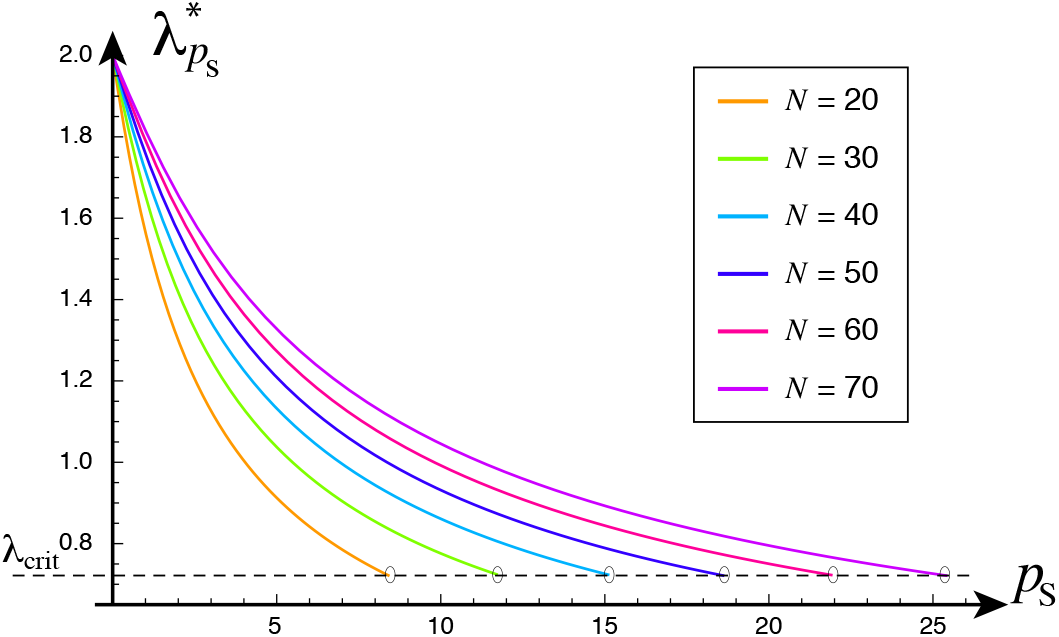
The impact of connectivity on the critical network toxic seeding – for a network consisting of one node with *N* neighbors. Plot of clearance *λ* versus toxic seeding *p_S_*. *ρ* = 6.4 × 10^−2^, while other parameters as in Figure 5.

These conceptual observations may be interpreted in the context of neurofibrillary tangle (NFT) staging [61, 81]. Indeed, (6) dictates that as toxic protein proliferates through neighboring regions, clearance is reduced below *λ*_crit_ and the formation of NFTs can result. Toxic infection will therefore generally take hold most rapidly in neighbors with lower clearance. Thus, the evolving distribution of clearance, and local toxic population growth, may affect the specific regional sequence of NFT staging. In addition to differences in NFT staging, patient-specific regional variations in clearance may also offer an explanation as to how extra-entorhinal seeding locations might emerge, as hypothesized in [81].

For instance, examining the relative values of the weighted graph Laplacian degree *d_ii_/d*_max_ (Figure 7), we note that the (right) entorhinal cortex (EC) is among the set of poorly supported regions. Thus, toxic τP seeds originating in the EC may tend to linger there. This observation is particularly interesting in the context of AD, consistent with observations from studies of τP staging [6, 19, 11] in AD, and motivates further research into modeling the region-specific balance and toxic protein load and of τP-related clearance mechanisms.

**Figure 7:**
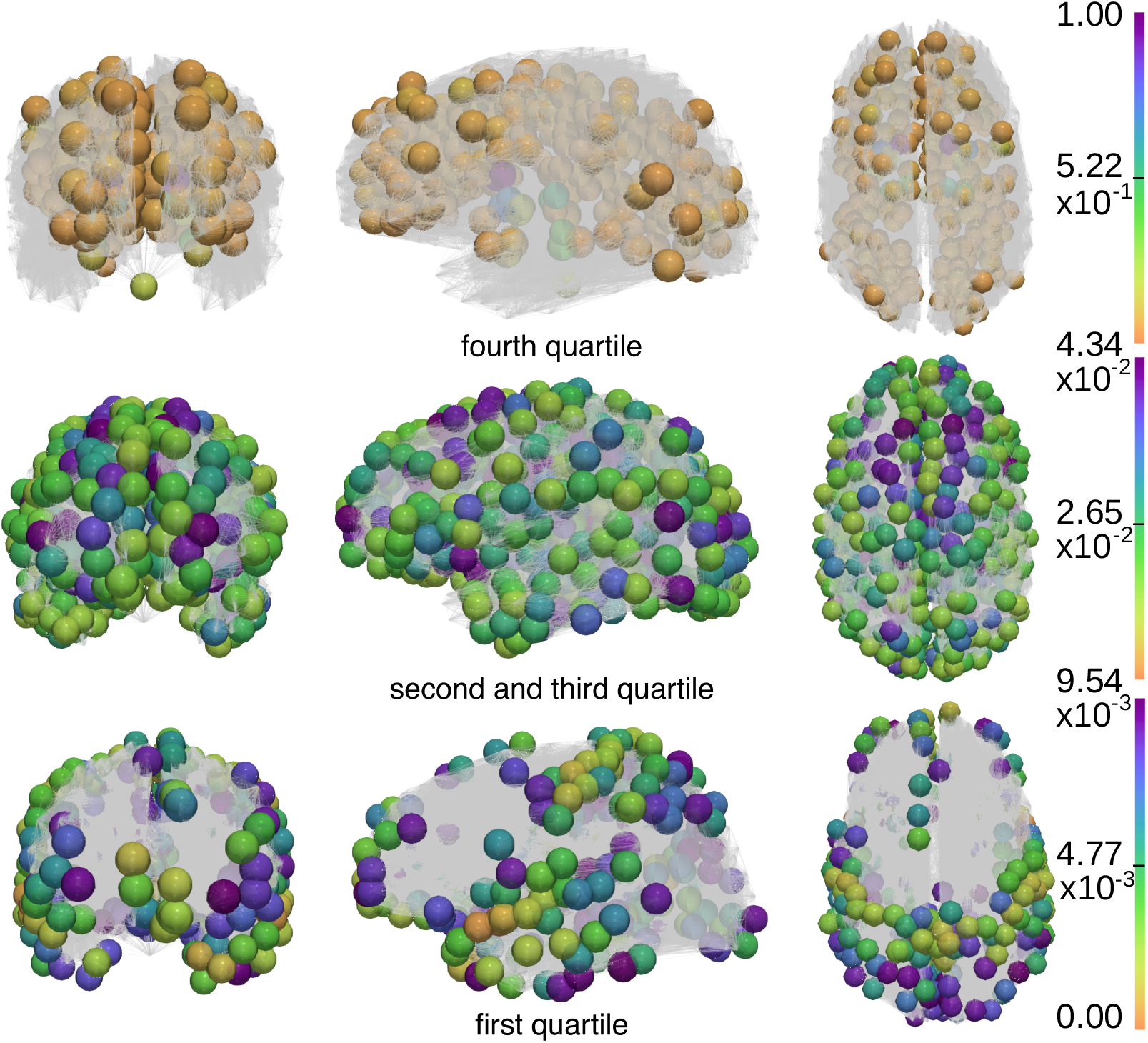
Relative total degree of each node in the connectome. Poorly supported regions (bottom row, first quartile values) with color scale 0 < *d_i_*/*d*_max_ ≤ 9.54 × 10^−3^, typical regions (middle row, second and third quartile values) with color scale 9.54 × 10^−3^ < *d_i_*/*d*_max_ ≤ 4.34 × 10^−2^ and well supported regions (top row, fourth quartile values) with color scale 4.34 × 10^−2^ < *d_i_*/*d*_max_ ≤ 1.

## 5 Dynamic brain clearance alters toxic protein progression

Finally, we investigate the full model (6) at the organ level by direct simulation. The simulations use a common set of model parameters (Table 1) where *ρ* and *α* were selected to produce maximal rates of toxic protein increase approximately on par with recent modeling studies employing AD imaging data [68, 67, 69]; *λ*_crit_ was chosen in line with experimental studies of aggregation kinetics [76]; 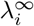 reflects the assumption that the regional clearance can reach a low but non-zero value; and *β_i_* was chosen to be one, which is consistent with a typical time-scale for disease progression of about 30 years. The computational results suggest that the distribution of clearance, throughout the various regions of the brain, may play a significant role in delaying disease onset, in producing the varied patterns of disease progression and can also serve as a mechanism that may explain some canonically studied AD subtypes.

**Table 1:**
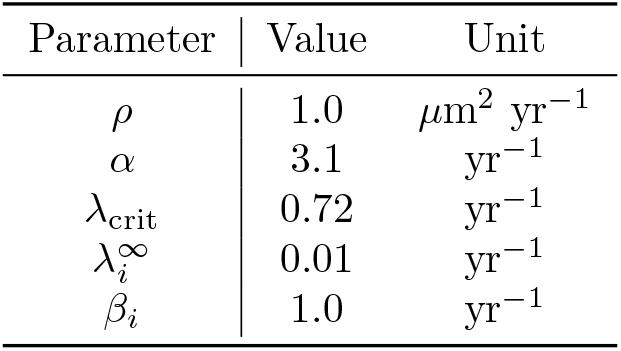
Parameter values used for organ-level simulations (Section 5).

### 5.1 Clearance delays disease onset and progression

We begin by mimicking a progression of τP in AD by placing an initial average toxic seeding *p*_0_ = 0.1 in each of the bilateral entorhinal cortices, alongside an initial clearance there of *λ*_0_ = *λ*^∞^. All other regions of the brain were initialized with *p*_0_ = 0.0 and *λ*_0_ = *γ*_sim_*λ*_crit_ (see Table 1). A series of simulations for seven different values of *γ*_sim_ ranging from 5% to 65% were performed. Note that the toxic protein concentration at each connectome node will saturate to the asymptotic value of *p*^*^ = 0.23 with this model set-up.

The computational results show that an increase in the level of healthy, homeostatic clearance delays the onset and progression of neurodegeneration (Figure 8). The lowest and highest initial clearance rates tested (*λ*_0_ = 5% *λ*_crit_, *λ*_0_ = 65% *λ*_crit_) yield onset times of *t* = 37.6 and *t* = 93.2 years, respectively, corresponding to a relative increase of 148%. A nonlinearly increasing relationship was noted between the initial clearance and onset time across the simulations (Figure 8, top right). Improved brain clearance, especially before neurodegenerative onset, may thus have significant benefits to brain health.

**Figure 8:**
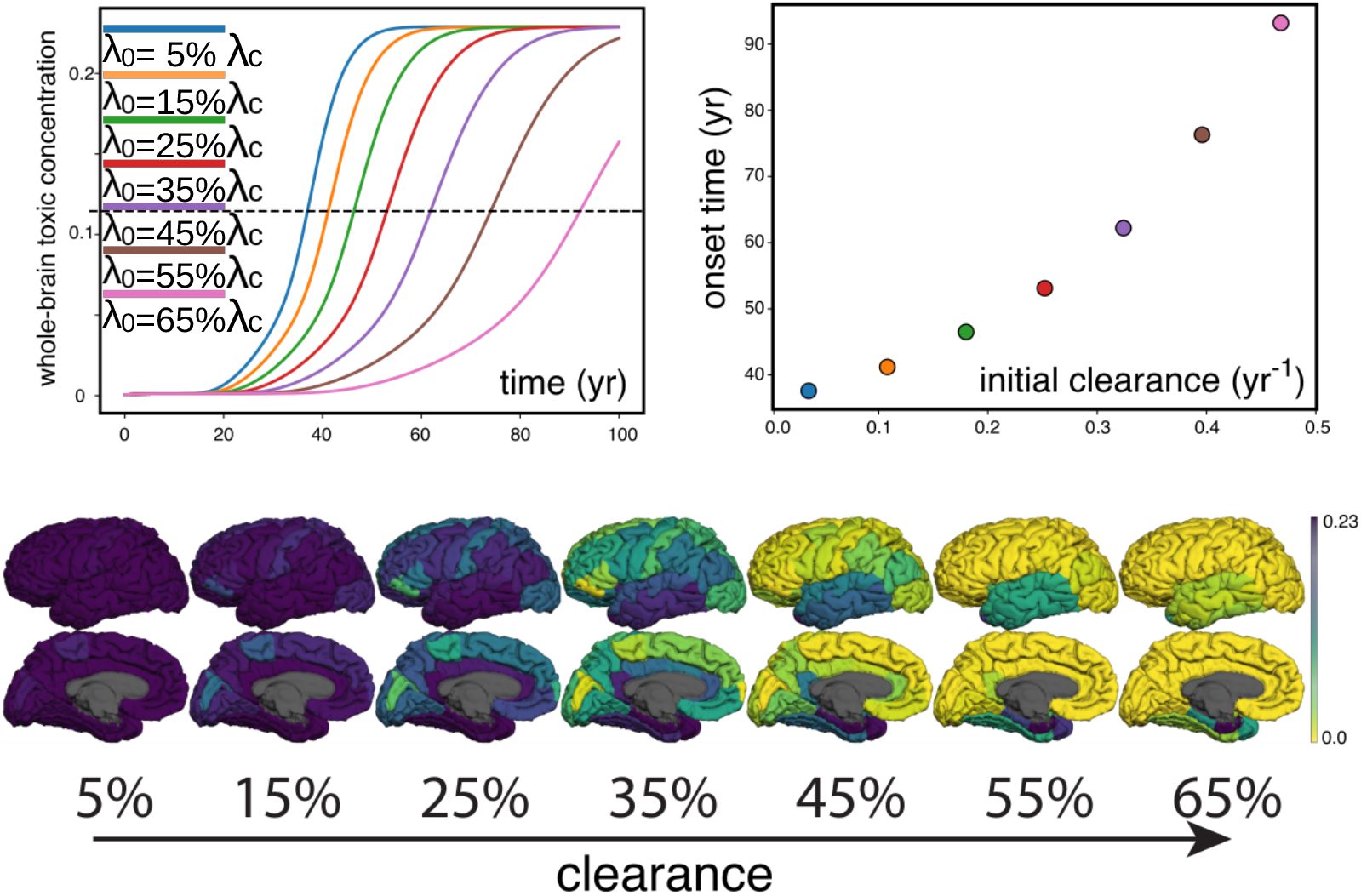
Neurodegenerative progression and time of onset. Top-left: Whole-brain average toxic load versus time for different levels of initial clearance. Top-right: the onset time (in years) at which 50% of the toxic protein saturation is reached versus the initial clearance level. Bottom: For each initial clearance, we show the toxic protein concentration at time *t* = 53.1 (median onset time, corresponding to *λ*_0_ = 35% *λ*_crit_).

Regional toxic burden was also seen to vary with initial clearance (Figure 8, bottom). In particular, and in line with the whole-brain average concentrations, the toxic protein load in any fixed region, at the median arrival time of *t* = 53.1, decreases with increasing initial clearance. In addition, a higher initial clearance is associated with a limbic and temporal predominance. Decreasing values of initial clearance are associated with increased temporal, parietal and frontal burdens. Furthermore, the observed progression of τP burden, as a function of initial clearance from right to left in Figure 8 (bottom row), is similar to experimentally observed τP NFT progression [38, Fig. 1f].

### 5.2 Spatial variations in clearance alter toxic protein progression

Neuroimaging studies suggest that the clearance capacity within the brain varies regionally and may be altered by age-related factors [74]. Assessments of perfusion [90], ubiquitination [70, 71, 42] and perivascular CSF circulation [66, 21, 22] point to specific regional differences as well as temporal variation in the major modes of brain clearance [74]. Here, we will demonstrate that regional variations in initial clearance can cause striking differences in the propagation of protein pathology. Since the connectivity of brain structural connectomes is complex, we first use an idealized geometry in order to demonstrate that clearance perturbs the flow of protein pathology by producing a toxic front that moves orthogonal to the gradient of the clearance field. We next extend these ideas to the connectome in Section 5.3.

We first consider four test cases defined over a uniform lattice of the unit square comprised of 100 equally spaced grid points in each direction. Each test case is initialized with the same toxic seeding concentration, *p*_0_ = 0.01, at the origin node (0, 0) with initial clearance set to *λ*^∞^. At all other nodes *p_i_*(0) = 0. We define four different distributions for *λ_i_*(0): labelled as <u>uniform</u>, <u>Gaussian</u>, <u>diagonal</u> and <u>linear</u>, shown in Figure 9 and described further below. For the uniform case, we set *λ_i_*(0) = *λ*^∞^, *i*. For the Gaussian case, the initial clearance is set according to a Gaussian distribution with *λ_i_*(0) = *λ*^∞^ + |*γ*|, where *γ* is a real value selected from a normal distribution with with mean *μ* = *λ*^∞^ and a standard deviation of *σ* = *λ*^∞^/2. For the diagonal case, we set *λ_i_*(0) = *λ*^∞^ in a central diagonal band and *λ_i_*(0) = 3*λ*^∞^ otherwise. The linear case sets an initial clearance field by *λ_i_*(0) = *λ*^∞^ at all nodes along the *y* = 0 line, and increasing linearly towards a maximum of *λ_i_*(0) = 3*λ*^∞^ along the line *y* = 1. Other parameters are taken from Table 1 and we set the total simulation time to be *t* = 100.

**Figure 9:**
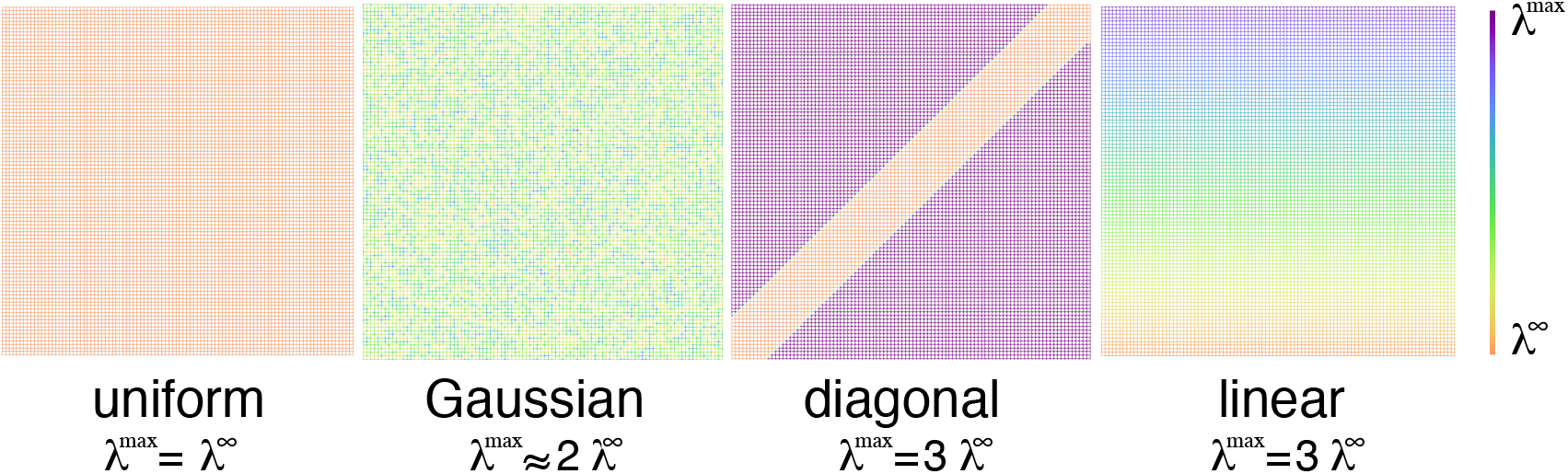
Four different idealized distributions of initial clearance defined on a uniform lattice of the unit square.

The corresponding simulated toxic protein progressions are shown in Figure 10. To increase the visibility of the flow front, toxic protein concentrations near zero are transparent while those coinciding with the onset value, determined by half of the maximal saturation value of *p_i_* = 0.23 (purple), are opaque. The toxic propagation first develops orthogonal to the clearance gradient, if such exists, subject to the underlying graph topology. With the uniform initial distribution of clearance, there is no gradient and the toxic front is constrained only by the topological connectivity of the graph. Similarly, the (discrete) gradient of the initial Gaussian clearance field is also Gaussian with *μ* = 0, and 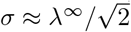. Hence, a clear sense of orthogonality is lacking also in this second test case, and we observe that the toxic front spreads in all connected directions. The diagonal test has a sharp gradient in the initial clearance field which is clearly reflected in the toxic protein propagation pathway. Finally, the fourth test case exhibits a constant clearance gradient oriented along the *y*-axis and the resulting toxic front advances first along the *x*-axis before propagating upwards.

**Figure 10:**
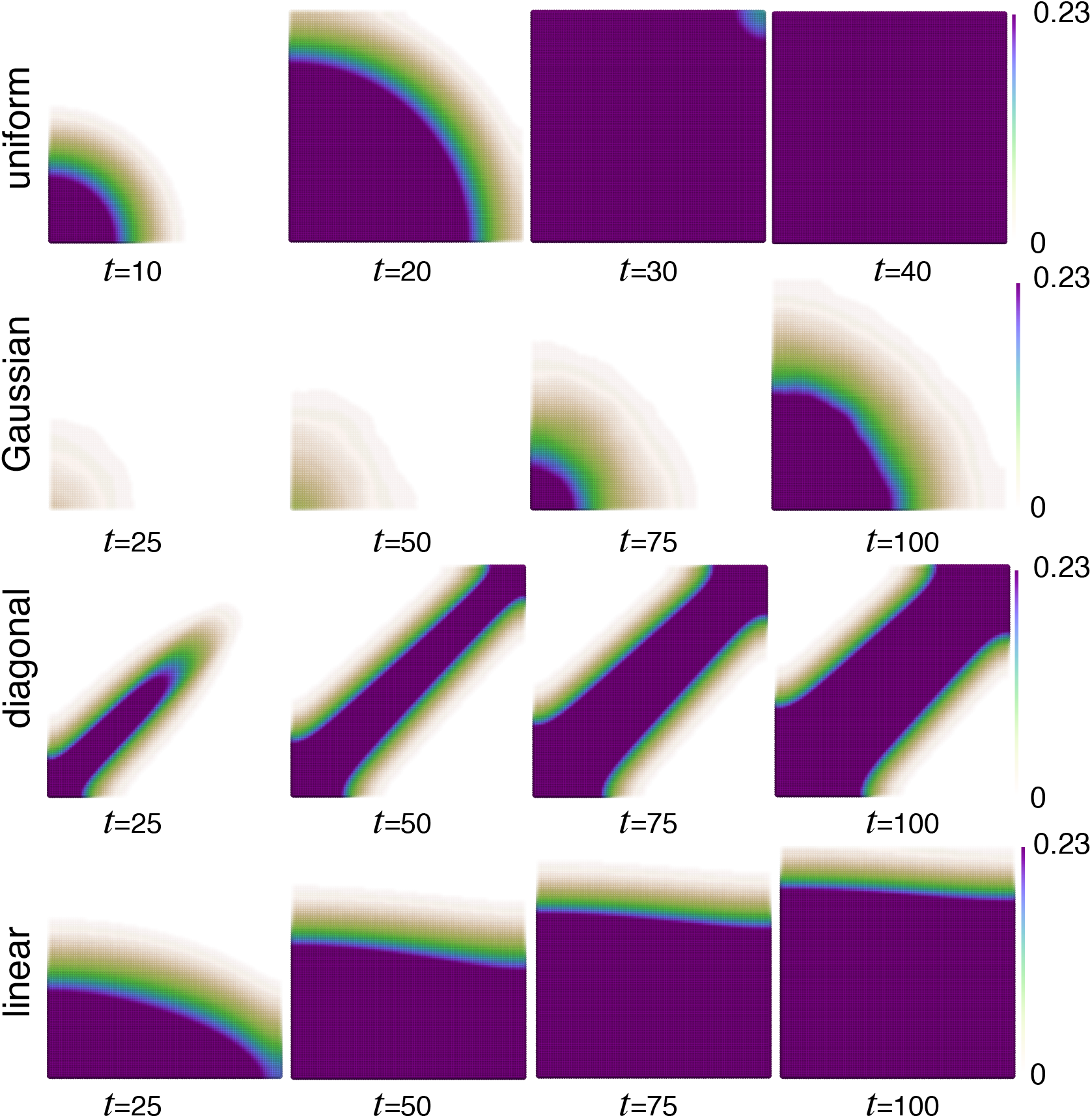
Simulated toxic protein progression at various simulation times for different initial clearance distributions corresponding to Figure 9. The maximal coloring (purple) corresponds to half the saturation value.

These results both echo and extend the observations of Section 5.1. The pattern with uniform or Gaussian initial distributions are similar, but with a smaller time scale in the Gaussian case due to a lower mean clearance. The case with diagonal or linear initial distributions extend this perspective by demonstrating that the direction of the initial clearance gradient can significantly alter the evolution of pathology and that pathology spreads most rapidly in the direction orthogonal to the gradient of clearance. Overall, these results strongly suggest that variations in brain clearance may significantly alter the patterns of toxic protein deposition in neurodegenerative diseases and may have further implications for the various trajectories [81] of τP deposition related to Alzheimer’s disease.

### 5.3 Clearance variation may promote AD subtypes

Recent studies have used τP progression to define notions of AD subtypes and have assumed that these different pathologies stem from different seeding regions [23, 81]. Here, we test the alternative hypothesis that subtypes can arise from the same seeding region but with regional differences in clearance.

To define AD subtypes from histopathology, postmortem NFT distributions of nearly two thousand patients of Braak stage V or later were collected in previous studies [47, 86, 37]. For each brain, these studies counted NFTs in the hippocampus (HP) and in the association cortex (ASC), where the latter was defined by superior temporal, middle frontal and inferior parietal regions (Figure 11). The ratio of HP to ASC NFTs (scores) was then computed, and the overall cohort distribution of values was determined [47]. The AD subtype classification is as follows:

- *Hippocampal sparing*: In this subtype, NFTs invade the association cortex more than the hippocampal region. It is defined by scores of less than the 25th percentile in the cohort distribution.
- *Typical AD*: Here, NFTs invade both the association cortex and the hippocampal region and is defined by scores between the 25th and 75th percentiles in the cohort distribution.
- *Limbic predominant*: NFTs invade the hippocampal region more than the association cortex and this subtype is defined by scores larger than the 75th percentile in the cohort distribution.

**Figure 11:**
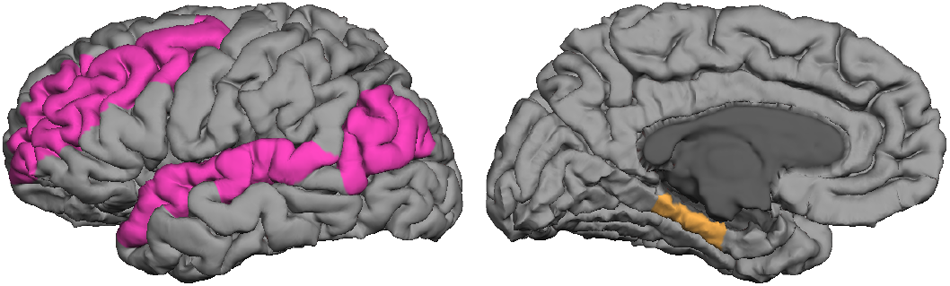
The classification of AD subtype depends on the number of NFTs in two particular regions, the association cortex (ASC) shown on the left and the hippocampus (HP) shown on the right. AD subtype with few NFTs in the HP regions compared to the ASC is called *hippocampus sparing*, whereas high values in the hippocampus regions compared to regions in the association cortex is called *limbic predominant*.

We will here demonstrate that simple variations in the initial distribution of clearance can elicit variations in the observed patterns of toxic protein progression and explain these AD subtypes.

To compare the effect of clearance on the distribution of NFTs, we follow previous studies [63, 75, 61, 67] and augment (6) with a measure of (nodal) NFT production, denoted by *q_i_*(*t*), reflecting damage accumulation following the arrival of toxic proteins. Given a toxic protein concentration *p_i_*(*t*), the (post-processed) NFT aggregation marker is defined as the solution of the damage equation:

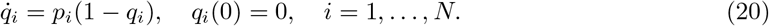

The variable *q_i_* is a local damage variable that increases from 0 to 1 as the disease progresses.

To measure the influence of clearance on the HP and ASC regions, we use the open-source *NetworkX* software package [27] to define *influential regions* for each. We consider the prevalence of the connection strengths between the nodes of a given composite ROI (either the HP or the ASC) and the nodes of its immediate neighbors; and by assessing how frequently a region appeared in shortest paths that originated in the EC and terminated in the HP or ASC composite ROIs (see Table 2).

**Table 2:** Influential regions for the composite hippocampal (HP) and association cortices (ASC) regions.

| Anatomical region | Influential regions |
| --- | --- |
| Hippocampus (HP) | Hippocampus, Insula, Thalamus |
| Association Cortices (ASC) | Banks of the superior temporal sulcus, Caudate,<br>Caudal middle frontal, Inferior parietal,<br>Pars triangularis, Precuneus, Putamen,<br>Rostral middle frontal, Superior temporal |

We investigate the progression of AD tauopathy by solving (6) and subsequently (20) for thirteen simulation scenarios. Each case uses the default model parameters (Table 1) along with initial clearance values for the HP influential region nodes set to *λ_i_*(0) = *λ*_crit_, and the initial conditions (*p_i_, λ_i_*(0)) = (0.01, *λ*^∞^) in the bilateral entorhinal cortices. The other initial clearance values *λ_i_*(0) are defined as follows. We consider 13 equispaced values of *M* ∈ [0.7, 1]. For each *M*, the initial clearance in the ASC influential regions is set to *λ*(0) = *Mλ*_crit_. The initial clearance in all other regions is set to *λ_i_*(0) = 1.8*λ*_crit_. Next, from the evolution of the damage *q_i_* in each region, we obtain *q*_HP_ as the average of (20) over the nodes of the hippocampal region and analogously for *q*_ASC_. For each simulation, we record the <u>onset time</u> at which either *q*_HP_ or *q*_ASC_ first reach 50%. Finally, each simulation result is classified according to the postmortem methodology of determining the ratio of *q*_HP_/*q*_ASC_ and assigning the subtype category based on its quartile range [47, 37, 86].

Our computations reveal that higher initial clearances in the ASC influential regions (*M* ∈ [0.925, 1.0]) produce a limbic predominant subtype, medium values (*M* ∈ [0.85, 0.9]) yield typical AD, while lower values (*M* ∈ [0.7, 0.825]) result in a hippocampal sparing subtype. Figure 12 shows four representative examples along the simulated type spectrum. Moreover, we find that the onset times cluster into similar groupings (Figure 13). These results reproduce the medical research observation that the hippocampal sparing variant reaches onset before typical AD, which itself reaches onset before the limbic predominant variant [47, 37, 86]. We conclude that regional variations in brain clearance may explain AD subtypes – an observation that motivates further studies in this direction.

**Figure 12:**
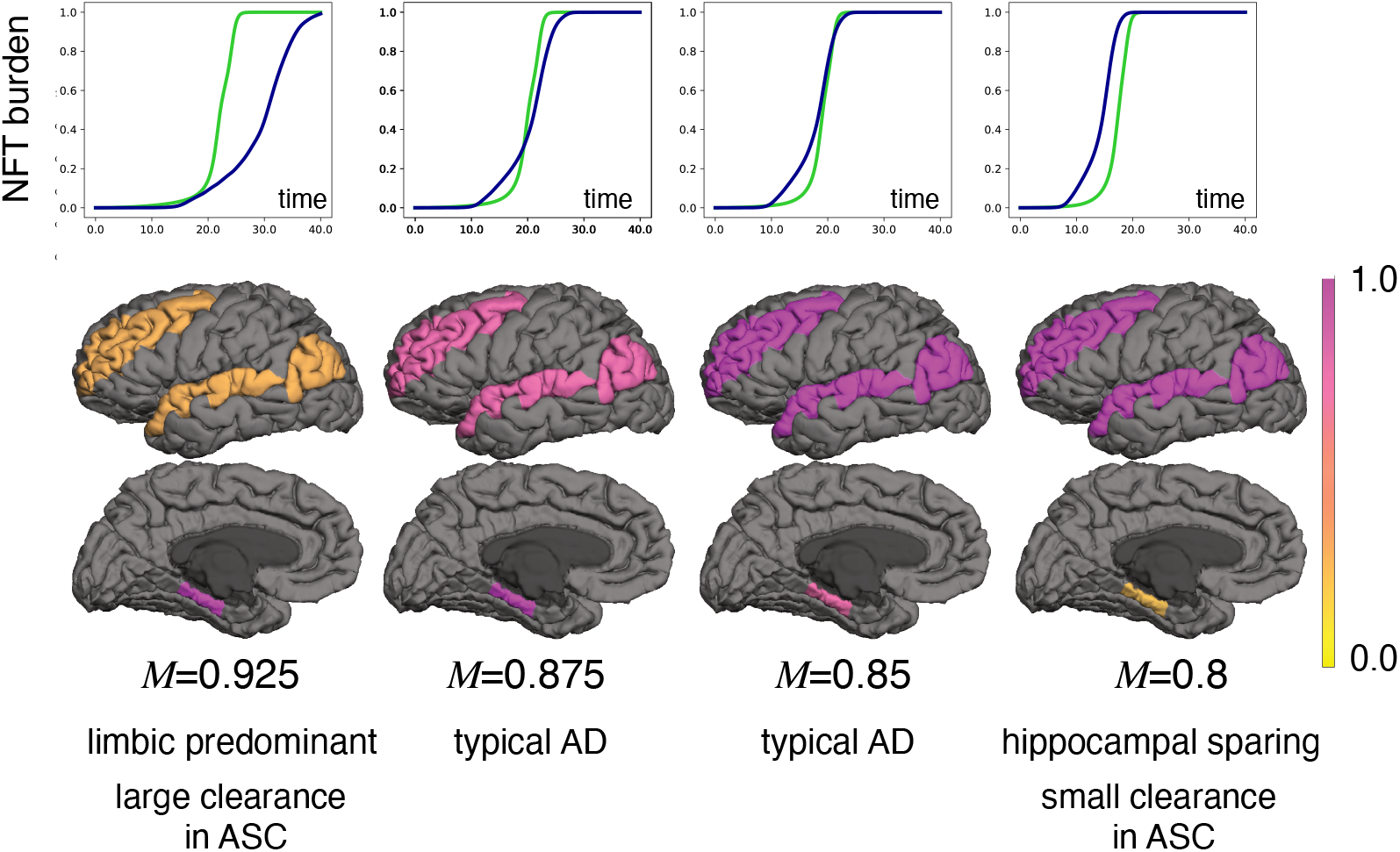
Simulated AD subtypes with average NFT aggregation marker (top row) in the HP (*q*_HP_ in green) and ASC (*q*_ASC_ in blue). For each *M*, we compute the time of onset which is defined as the first time that either *q*_HP_ or *q*_ASC_ reaches 1/2. The value of the NFT aggregation marker (20) in the ASC (middle row) and HP (bottom row) are then plotted, at time of onset, for each of the corresponding NFT aggregation plots (top row).

**Figure 13:**
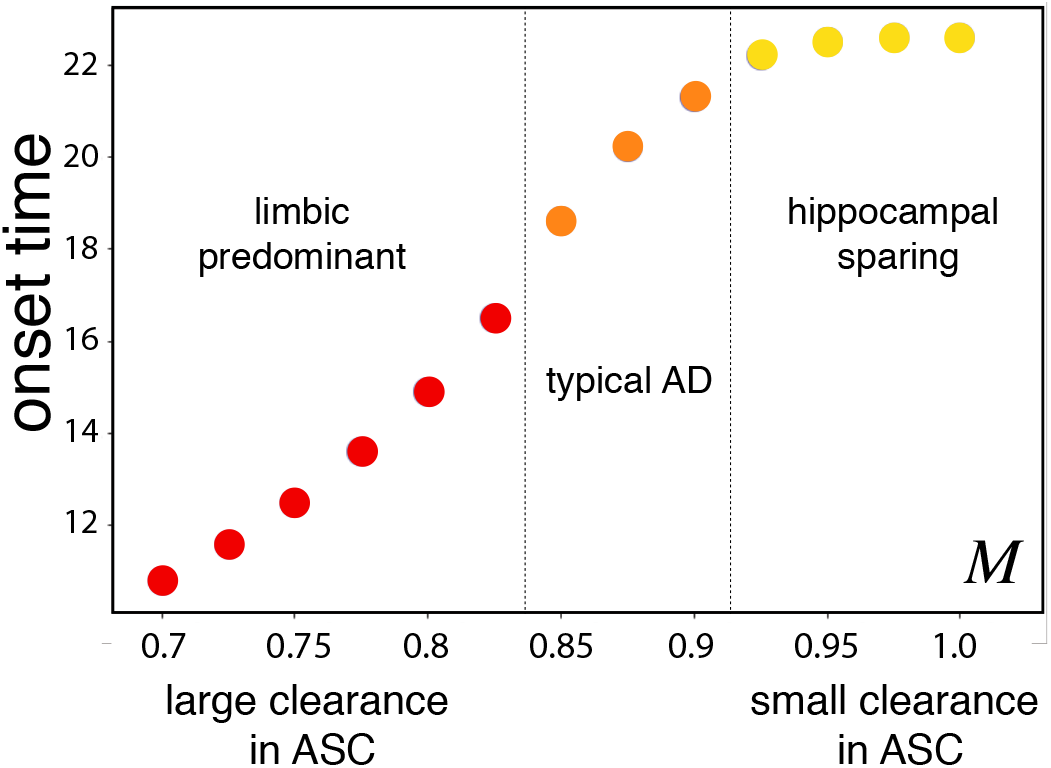
Simulated time of onset for different simulation scenarios corresponding to different initial clearance modifier values *M*, grouped by AD subtypes.

## 6 Conclusion

Mathematical models that make use of an underlying brain network have been used to study patient data [34, 82, 68, 81, 67] and to examine the implications of mechanisms, largely observed in or hypothesized from animal model experiments, in the progression of neurodegeneration over human brain networks [62, 63, 54, 24, 25, 75]. A constant brain clearance parameter has been included in some of these models [24, 25, 34, 75, 81, 82]. However, given the fundamental role that brain clearance systems are thought to play in neurodegenerative diseases [72, 74, 88, 64, 2, 44, 50, 31], and that brain clearance is now thought to deteriorate in the presence of toxic proteins [4, 9, 10, 31, 50], the careful study of the implications of an evolving brain clearance is all but overdue.

Our model is the first of its kind to study the coupled progression of clearance and protein pathology. A previous network neurodegeneration model has suggested that brain clearance may explain why Alzheimer’s disease is a generally considered to be a secondary tauopathy [75] but this model fails to consider how brain clearance may be altered by pathological proteins and thus, ultimately, fails to account for the important role that clearance may play in shaping brain vulnerability to toxic invasion and in explaining neurodegenerative subtypes. Conversely other models have used mathematical expressions with constant, regional parameters [34, 82, 81], including clearance, and have fitted these parameters to large quantities of patient neuroimaging data. These types of studies are invaluable for finding patterns in human neuroimaging data that are anticipated, or hypothesized, from animal model experimental evidence and can give a good idea as to how a patient may progress based on previous longitudinal, large cohort data studies. However, fitting large volumes of data is necessarily tied to a particular set of data and offers only a little in the way of a principled mechanistic understanding of the implications of couplings between disease progression and a patient’s evolving in vivo environment.

Our model begins by codifying the observation that clearance directly affects the dynamics of a misfolded protein species within the brain [76] by inducing an instability in the microscale aggregation kinetics. We then start with a healthy brain that has sufficient clearance to eliminate an amount of toxic proteins and maintain a stable state of homeostasis. However, as toxic proteins increase, brain clearance system become increasingly damaged; the brain is subject to full invasion as reduced clearance leads to an unstable, aggregation prone state and the prion-like reproduction and proliferation of toxic, neurodegenerative proteins dominates. In the absence of a model including an evolving clearance, the regions that have sufficient clearance will always be protected. Our model of coupled clearance and neurodegeneration provides insights for future research. Our analysis suggests that the brain may exhibit clearance-dependent regional homeostasis and that the topology of the brain may provide resilience against a toxic protein infection taking hold. Our simulations, motivated by the progression of τP in AD, further suggest that brain clearance systems are a likely therapeutic target for future drug development research. In particular, we show that increasing clearance in a healthy brain may be instrumental in significantly delaying AD onset, that the progression of toxic pathology depends on regional clearance levels and that patient-specific distribution of brain clearance may play a key role in the manifestation of AD subtypes.

Our study presents with a number of limitations and opportunities for continued research. First, the enormous complexity of the brain is clearly not fully describable by a network graph; many local, and global, effects must be ignored in order to make this geometrical simplification. For instance, network dynamical systems models reflect average changes in regional variables (vertices) that depend strongly (edges) on the state of a select group of neighbors. Thus, regional dynamics are significantly homogenized, local spatial resolution is lost and some quantities of interest may be unrecoverable. Second, a brain connectome graph (Figure 3) does not represent structures that allow for the multiphysics mathematical modeling of many important phenomena, with implications for neurodegenerative diseases, such as cerebrospinal fluid circulation, vascular stenosis, cerebral autoregulation, stroke, oxygen transport, endothelial dysfunction or the regional effects of traumatic injury, among others. Third, neurodegenerative diseases often involve fundamental interactions between more than one brain protein [3, 33, 65, 80] and multiple clearance mechanisms [57, 74, 79]. However, our model only includes one toxic protein species and combines distinct brain clearance systems into a single (regional) term; extensions of the single compartment protein and clearance model may provide additional insights. Finally, our model has yet to be assessed using regional clearance or toxic concentration values garnered from neuroimaging data sets, or derived directly from an experimental setting.

Despite the limitations presented by the current study, we have advanced the first model that relates an instability in protein aggregation, the brain’s dynamic clearance capacity and the prion-like replication and proliferation of misfolded proteins in neurodegenerative diseases. The analysis of the model has revealed striking insights into how clearance deficits may engender disease proliferation, the notion of region-specific critical seeding, regional vulnerability due to variations in connectivity, the importance of regional clearance in delaying the onset of neurodegeneration and the implications for clearance in determining neuropathologically defined Alzheimer’s disease subtypes. Our mathematical evidence supports a growing hypothesis of the medical community: brain clearance plays an important role in the etiology and progression of neurodegenerative diseases.

## Acknowledgement

This publication is based on work supported by the EPSRC Centre For Doctoral Training in Industrially Focused Mathematical Modelling (EP/L015803/1) in collaboration with Simula Research Laboratory. The work of A. Goriely was supported by the Engineering and Physical Sciences Research Council (EPSRC) grant EP/R020205/1. M. E. Rognes has received funding from the European Research Council (ERC) under the European Union’s Horizon 2020 research and innovation programme under grant agreement 714892. The work of G.S. Brennan was partially supported by the EPSRC InFOMM grant EP/L015803/1. The work of T.B. Thompson was supported partially by the John Fell Oxford University Press Research Fund grant BKD00160 and partially by the EPSRC grant EP/R020205/1 to AG. The work of H. Oliveri was funded by the EPSRC grant EP/R020205/1 to AG.

